# dreampy: Pseudobulk mixed-model differential expression for single-cell RNA-seq in Python

**DOI:** 10.64898/2026.03.21.713408

**Authors:** Steven B. Wells, Hamna Shahnawaz, Joanne L. Jones

## Abstract

dreampy is a Python implementation of the R dreamlet framework for pseudobulk differential expression analysis of single-cell RNA-seq data. dreamlet combines voom precision-weighted linear mixed models with empirical Bayes moderation to handle batch effects, repeated measures, and other hierarchical structure in multi-donor studies, but exists entirely within the R/Bioconductor ecosystem. dreampy reproduces this pipeline natively in Python, integrating with AnnData and the scverse ecosystem.

## Introduction

Large-scale single-cell and single-nucleus RNA-seq studies now routinely profile hundreds of thousands to millions of cells across hundreds of donors, enabling systematic investigation of how gene expression varies with disease, treatment, and other phenotypes at cell-type resolution. A central analytical challenge in these studies is differential expression (DE) testing: identifying genes whose expression differs between conditions while properly accounting for the hierarchical structure of the data. In these datasets multiple cells are observed per donor, donors may contribute samples across batches or tissues, and technical variation is introduced at multiple levels.

Early approaches to single-cell DE analysis applied cell-level statistical tests, treating each cell as an independent observation. Multiple groups subsequently demonstrated that this inflates false positive rates dramatically, as cells from the same donor are not independent — a problem known as pseudoreplication (Crowell et al. 2020; Squair et al. 2021; Zimmerman et al. 2021). Therefore, the field has converged on pseudobulk aggregation as the preferred remedy: summing counts across cells within each donor–cell type combination to produce one observation per biological replicate, then applying bulk RNA-seq statistical frameworks to the aggregated counts. This approach respects the true unit of replication — the donor — and enables the use of mature, well-calibrated statistical methods.

Among pseudobulk DE frameworks, dreamlet (Hoffman et al., 2024) occupies a distinctive position. Rather than using negative binomial generalized linear models (as in edgeR or DESeq2), dreamlet builds on the limma-voom framework (Law et al. 2014; Smyth 2004), which transforms counts to log-counts-per-million with observation-level precision weights derived from an empirical mean–variance trend, then fits linear models to the transformed data. Critically, dreamlet extends this to linear mixed models with lme4 (Bates et al., 2015) estimating denominator degrees of freedom using either Satterthwaite approximation (lmerTest, Kuznetsova et al. 2017) or Kenward-Roger correction (pbkrtest, Halekoh and Højsgaard 2014), and applying empirical Bayes moderation via variancePartition (Hoffman and Schadt 2016) to the mixed-model output. This combination — voom precision weights, linear mixed models for complex experimental designs, and empirical Bayes shrinkage for borrowing strength across genes — provides a statistically principled framework that handles batch effects, repeated measures, and other random-effect structure naturally within the model, rather than requiring them to be absorbed into fixed effects or removed by upstream correction.

However, this statistical machinery exists entirely within the R/Bioconductor ecosystem. The core pipeline spans seven external packages (edgeR, limma, lme4, lmerTest, pbkrtest, fANCOVA, and variancePartition itself), each with its own dependencies, data structures, and interfaces. For researchers working primarily in Python — where scanpy (Wolf et al., 2018) and the broader scverse ecosystem (Virshup et al., 2023) have become the standard environment for single-cell analysis from preprocessing through cell type annotation — using dreamlet requires exporting data to R, running the analysis in a separate environment, and importing results back. This language-switching workflow is not merely inconvenient; it creates friction that discourages interactive exploration, complicates reproducibility, and makes it difficult to integrate DE results with downstream Python-based analyses.

Recent efforts have begun to address the broader R-to-Python gap for differential expression. PyDESeq2 (Muzellec et al., 2023) reimplements the DESeq2 framework in Python but supports only fixed-effects models for bulk RNA-seq, with no mixed-model capability. edgePython (Pachter, 2026) provides a broad port of edgeR and extends it with a negative binomial–gamma mixed model following the NEBULA-LN approach (He et al., 2021), applying empirical Bayes shrinkage to cell-level dispersions — a distinct statistical framework from the voom-weighted linear mixed models used by dreamlet. InMoose (Colange et al., 2025) ports limma, edgeR, and DESeq2 to Python but does not include mixed-model support. None of these tools provide the limma-voom pipeline with linear mixed model support, Satterthwaite or Kenward-Roger degrees of freedom, and empirical Bayes moderation of residual variances that characterize the dreamlet framework.

Here we present dreampy, a native Python implementation of the dreamlet pseudobulk mixed-model workflow. dreampy reimplements the full pipeline — from pseudobulk aggregation and TMM normalization through voom precision weighting, mixed-model fitting, and empirical Bayes moderation — using Python’s scientific computing stack (NumPy, SciPy, pandas) and integrating directly with AnnData, the standard data structure in the scverse ecosystem. The package supports both fixed-effect and random-effect formulas, dispatching to ordinary least squares or restricted maximum likelihood estimation as appropriate, and exposes every pipeline stage as an individual, inspectable function call — in contrast to R dreamlet, which bundles most stages behind two user-facing entry points (processAssays() and dreamlet()).

We validate dreampy against R dreamlet on two published datasets, demonstrating Pearson correlations up to r = 0.9999997 on individual pipeline stages and passing 332 of 351 and 249 of 270 assay-level metric tests at a correlation floor of r ⩾ 0.999. As a biological application, we reanalyze a lupus cohort (Perez et al., 2022) using a mixed-effects model that recovers control donors excluded from the original analysis due to batch aliasing in their fixed-effects framework, demonstrating both increased statistical power and robust detection of the canonical interferon-stimulated gene signature across eight immune cell types.

### Implementation

dreampy reimplements the complete dreamlet pseudobulk mixed-model pipeline as a sequence of nine composable Python functions, each corresponding to a distinct statistical operation (Figure 1). This explicit decomposition contrasts with R dreamlet, where six of the nine stages are bundled behind two user-facing entry points — processAssays() (aggregation through voom weighting) and dreamlet() (model fitting through results extraction) — with the underlying statistical engines distributed across seven external packages. dreampy’s flat architecture gives users direct access to every intermediate result, facilitating inspection, debugging, and customization of any pipeline stage.

**Figure 1:**
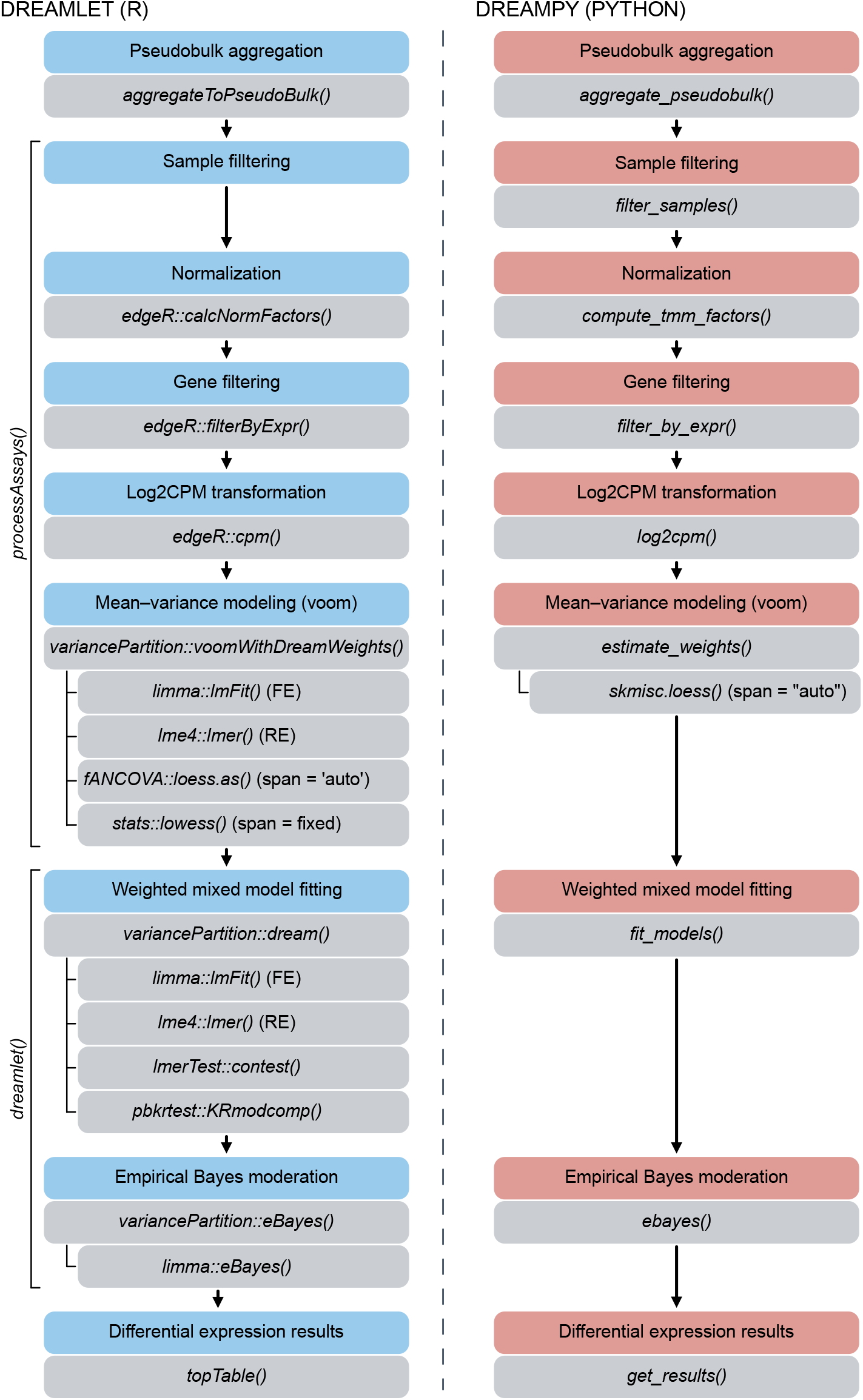
Pipeline architecture of R dreamlet and dreampy. Side-by-side comparison of the R dreamlet (left) and dreampy (right) pipelines. R dreamlet bundles six pipeline stages behind two user-facing entry points — processAssays() (pseudobulk aggregation through voom precision weighting) and dreamlet() (weighted model fitting through differential expression results) — with statistical operations distributed across seven external packages (edgeR, limma, lme4, lmerTest, pbkrtest, fANCOVA, variancePartition). dreampy exposes each stage as an individual function call, from aggregate_pseudobulk() through get_results(), with formula-based dispatch to fixed-effect (OLS) or random-effect (REML via BOBYQA) fitting paths. Both pipelines produce identical statistical outputs; dreampy integrates natively with AnnData and the Python scverse ecosystem.

#### Pipeline stages

The pipeline proceeds as follows: aggregate_pseudobulk() sums raw counts across cells within each donor–cell type combination, producing an AnnData object with pseudobulk profiles as observations and cell types stored as an assay label. filter_samples() removes pseudobulk samples falling below a minimum cell count threshold and drops cell types with too few remaining samples. compute_tmm_factors() estimates trimmed mean of M-values normalization factors (Robinson & Oshlack, 2010) from the pseudobulk count matrix. filter_by_expr() removes lowly expressed genes using an expression-level filter that accounts for library size and experimental design, reimplementing edgeR’s filterByExpr() function, which codifies the filtering strategy described in Chen et al. (2016). log2cpm() transforms counts to log2 counts per million with a constant prior count of 0.5 — matching limma-voom’s behavior, where the prior is not scaled by library size. estimate_weights() performs the voom mean–variance modeling step. For each gene, an initial linear model (fixed effects) or linear mixed model (random effects present) is fit to the log2 CPM values without weights, yielding per-gene residual standard deviations and fitted values. A nonparametric smooth is then fitted to the square-root residual standard deviation as a function of mean log-count, and precision weights are derived as the inverse fourth power of the estimated trend evaluated at each observation’s fitted value. Two smoothing strategies are supported, matching the behavior of R dreamlet: when span=‘auto’, a local polynomial regression (loess) is fit with the smoothing parameter selected by minimizing the bias-corrected Akaike information criterion (AICc; Hurvich et al. 1998), implemented via scikit-misc’s loess function and matching the behavior of R’s fANCOVA::loess.as(); when span is set to a fixed value (e.g., 0.5), a lowess smooth is used instead, matching R’s stats::lowess(). Cross-language validation uses a fixed span=0.5 (the lowess path) because the automatic span selection can produce small differences between the R and Python loess implementations, and fixing the span isolates validation of the statistical pipeline from smoother-selection variability.

fit_models() fits the full precision-weighted linear model to each gene in parallel. When the formula contains no random effects, ordinary weighted least squares is applied. When random effects are present, a weighted linear mixed model is fit by optimizing the profiled REML (or ML) deviance over the variance component parameters *θ*, using the BOBYQA derivative-free optimizer (Powell, 2009) as implemented in Py-BOBYQA (Cartis et al., 2019). For each gene, the Satterthwaite approximation (Satterthwaite, 1946) is used to estimate denominator degrees of freedom for each fixed-effect coefficient, providing per-coefficient degrees of freedom that account for the variance component uncertainty. Kenward-Roger degrees of freedom (Halekoh & Højsgaard, 2014) are also implemented and validated but are not used in the default pipeline due to their substantially higher computational cost. Following model fitting, ebayes() applies empirical Bayes moderation (Smyth, 2004), shrinking gene-wise residual variances toward a common prior to stabilize inference, particularly for genes with few residual degrees of freedom. The moderated t-statistics are computed using per-coefficient Satterthwaite degrees of freedom for hypothesis testing, while the prior variance estimation itself uses the trace-based residual degrees of freedom — matching the bookkeeping in variancePartition’s eBayes(). Finally, get_results() extracts a per-gene results table with coefficients, moderated t-statistics, p-values, Benjamini-Hochberg adjusted p-values, and log-odds of differential expression.

#### Design decisions

Several design choices distinguish dreampy from the R implementation beyond the language difference.

#### Cold start initialization

R dreamlet warm-starts the optimizer for each gene using the previous gene’s converged variance components within each parallel worker. This introduces a gene-order dependency: results can differ depending on how genes are distributed across workers. dreampy instead computes independent starting values for each gene using a method-of-moments heuristic applied to the random-effect design matrix and the response. This cold-start approach is deterministic regardless of gene ordering or parallelization strategy, at the cost of occasionally converging to a different local optimum on flat or multimodal likelihood surfaces.

#### REML defaults

R dreamlet uses maximum likelihood (ML) for the voom weight estimation step and REML for model fitting — an inconsistency likely inherited from limma-voom, which predates mixed-model support and has no REML concept. dreampy employs REML for both stages, providing a uniform default that is more appropriate for variance component estimation in the moderate sample sizes typical of pseudobulk designs. All cross-language validation uses R’s ML/REML settings to demonstrate numerical concordance under identical conditions.

#### Explicit collinearity handling

When random-effect terms are perfectly collinear (e.g., a donor who appears in only one batch, creating a rank-deficient random-effects design matrix), dreampy detects and drops the redundant term before fitting, producing a reduced but well-identified model. R dreamlet retains the full parameterization and relies on the optimizer to handle the resulting near-singularity via theta boundary behavior. Both approaches yield concordant differential expression calls on shared genes, but dreampy’s explicit handling avoids convergence failures on genes where R’s optimizer struggles with the degenerate parameterization.

### Cross-language validation

We validated dreampy against R dreamlet using two published single-cell RNA-seq datasets that span different biological contexts, experimental designs, and sample sizes.

**Wells et al. (2025)** profiled T cell immune aging across donor and tissue site, yielding 13 T cell assays with 41 to 153 pseudobulk samples (donor × tissue), fit with the formula ∼*age*_*groups* + (1 | *donor*_*id*) + (1|*tissue*_*abbrev*). On the CD4 Treg assay (135 samples, 8,139 genes), dreampy and R dreamlet produce near-identical outputs at every pipeline stage, with Pearson correlations ranging from r = 0.9999997 (adjusted p-values) to r = 1.0000000 (TMM factors) and a maximum absolute difference of 5.33 × 10^−14^ on log2 CPM values (Figure 2a–g). Across all 13 assays, 332 of 351 per-assay metric tests pass at a correlation floor of r ⩾ 0.999. The 19 failures were concentrated in four small-sample T cell subtypes (CD8 MAIT, CD4 TEMRA, CD8 CM, CD8 TEMRA). Of these, 17 were correlation-floor violations on variance-component-sensitive quantities — theta estimates and downstream degrees of freedom — where optimizer boundary behavior on multimodal likelihood surfaces produces expected divergence between the two implementations’ starting-point strategies. The remaining two are floating-point tie-breaks in gene filtering at the CPM boundary (≈10^−15^ absolute difference), an inherent consequence of different floating-point operation ordering between R’s compiled C++ and NumPy.

**Figure 2:**
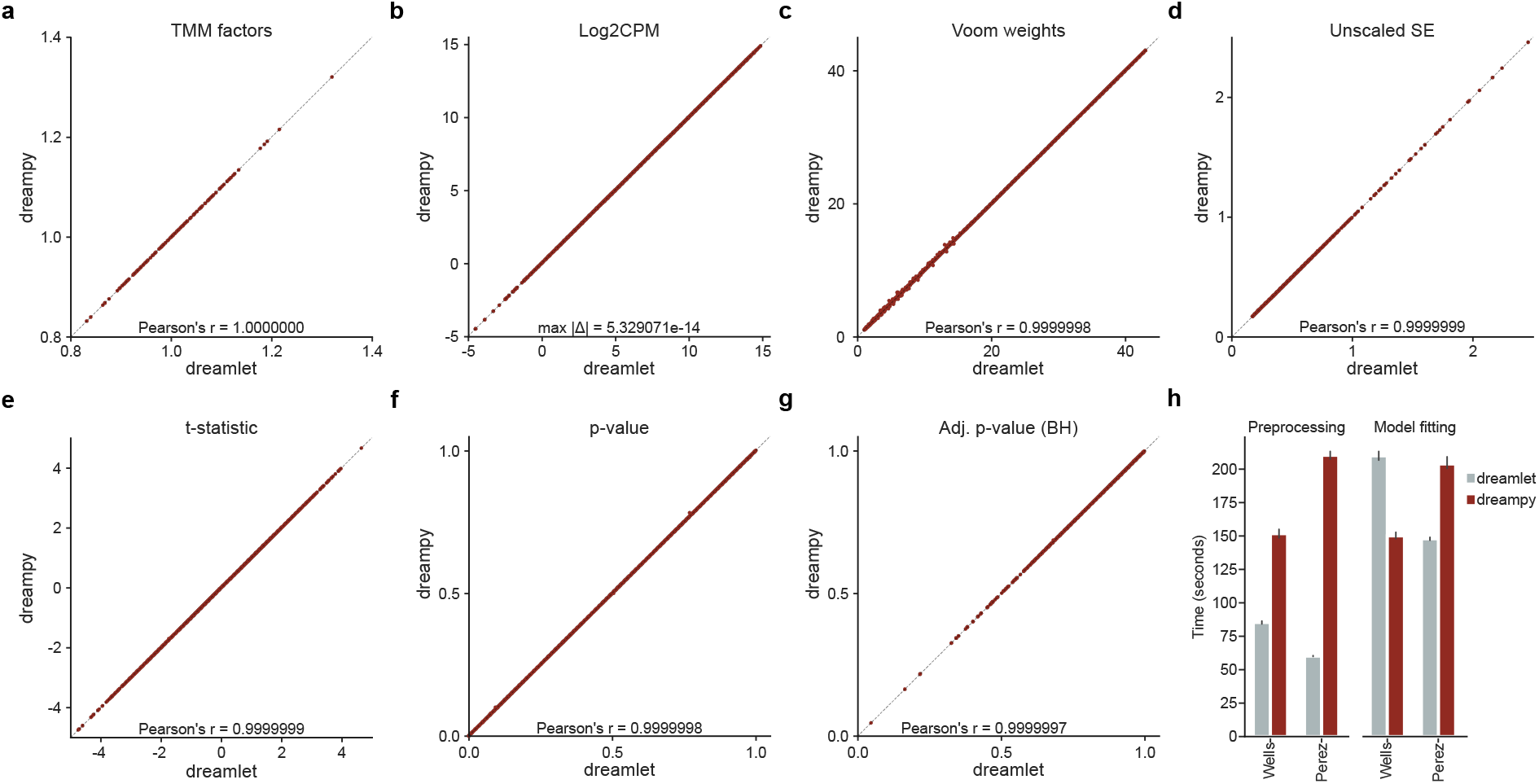
Cross-language validation and speed comparison. (a–g) Identity-line scatter plots comparing dreampy (y-axis) and R dreamlet (x-axis) outputs on the Wells et al. (2025) CD4 Treg assay (135 samples, 8,139 genes) across seven pipeline stages: (a) TMM normalization factors (Pearson r = 1.0000000), (b) log2 CPM values (maximum absolute difference ≈5.33 × 10^−14^), (c) voom precision weights (r = 0.9999998), (d) unscaled standard errors (r = 0.9999999), (e) moderated t-statistics (r = 0.9999999), (f) p-values (r = 0.9999998), (g) Benjamini-Hochberg adjusted p-values (r = 0.9999997). Dashed grey line indicates y = x. Correlations and summary statistics were computed on all observations; panels b and c display 10,000 randomly sampled points for visual clarity. (h) End-to-end wall-clock time comparison on the Wells and Perez datasets, split into preprocessing (pseudobulk aggregation through voom weighting) and model fitting (weighted models through eBayes). Both implementations used 16 cores on a MacBook Pro with an Apple M4 Max and 64 GB RAM running macOS 26.2. R dreamlet used MulticoreParam(16); dreampy used n_jobs=−1.

**Perez et al. (2022)** profiled lupus and healthy immune cells across 261 donors and 10 cell types, yielding 11 to 336 pseudobulk samples fit with ∼*sle* + (1 | *donor*_*id*) + (1 | *Processing*_*Cohort*). Of 270 assay-metric tests, 249 passed. The 21 failures included 15 from the plasmablast (PB) assay, where perfect collinearity between processing cohort and disease status forced dreampy’s collinearity detection to fit a structurally different one-random-effect model, and 6 from two small-population cell types (pDC, proliferating cells) attributable to the same optimizer boundary sensitivity observed in the Wells et al dataset. Despite the intermediate-quantity disagree-ment in the PB assay, both implementations identified identical differentially expressed genes with concordant direction and significance.

### Speed comparison

All benchmarks were performed on a MacBook Pro with an Apple M4 Max (16 cores: 12 performance, 4 efficiency) and 64 GB RAM, running macOS 26.2. Figure 2h compares end-to-end wall-clock times for both datasets, split into preprocessing (aggregation through voom weighting) and model fitting (weighted models through results), with both implementations using 16 cores (MulticoreParam(16) in R, n_jobs= −1 in dreampy). The results were mixed: dreampy was faster on Wells et al preprocessing but slower on model fitting, while the pattern was reversed on the Perez et al dataset. Neither implementation was consistently faster across datasets. dreampy’s cold-start strategy incurred per-gene overhead that is absent from R’s warm-start approach, while dreampy’s model construction avoided R’s per-gene S4 method dispatch overhead. The net effect depended on dataset characteristics — number of genes, sample size, number of random-effect levels, and the fraction of genes near variance component boundaries. Speed optimization, including warm-start options and a lean deviance-only fast path during optimization, is planned for v0.2.

## Results

### Reanalysis of a lupus cohort recovers excluded controls and strengthens the interferon signature

To demonstrate dreampy on a biological question where the mixed-model framework provides a concrete analytical advantage, we reanalyzed the Perez et al. (2022) lupus cohort (Methods).

The original Perez et al. analysis used edgeR with a fixed-effects model (∼*disease* + *Processing*_*Cohort* + *age*). As the processing cohort was modeled as a fixed effect, cohort 1 — which contained only ImmVar healthy controls — was perfectly aliased with disease status, forcing the authors to exclude all 50 ImmVar donors from case-control comparisons. This exclusion removed half of the healthy control group (99 to 49 donors), substantially reducing statistical power. Moreover, 21 of the excluded ImmVar donors bridged multiple processing cohorts, including 10 who were present in batch 3 alongside SLE cases, meaning the exclusion extended well beyond the aliased donors. A mixed-effects model avoids this aliasing by estimating cohort effects as random deviations from the population mean rather than as separate fixed-effect levels, allowing the ImmVar donors to contribute to inference.

We fit the model ∼ *sle* + (1 | *donor*_*id*) + (1 | *Processing*_*Cohort*), with the donor random effect absorbing repeated measures and the processing cohort random effect absorbing batch variation (Methods). The reanalysis was performed in two configurations: the full cohort (261 donors) and an ImmVar-excluded cohort (211 donors) that faithfully reproduced the Perez et al. exclusion. Comparing the two quantifies the statistical cost of the exclusion and serves as an internal sensitivity analysis.

Recovering the ImmVar controls substantially increased the number of detected DE genes across all eight cell types — for example, 3,905 versus 2,084 in classical monocytes, 3,643 versus 1,603 in CD4 T cells, and 2,841 versus 856 in CD8 T cells (Figure 3a). The pattern was consistent: the full cohort roughly doubles the number of DE genes (FDR < 0.05) for the major cell types.

**Figure 3:**
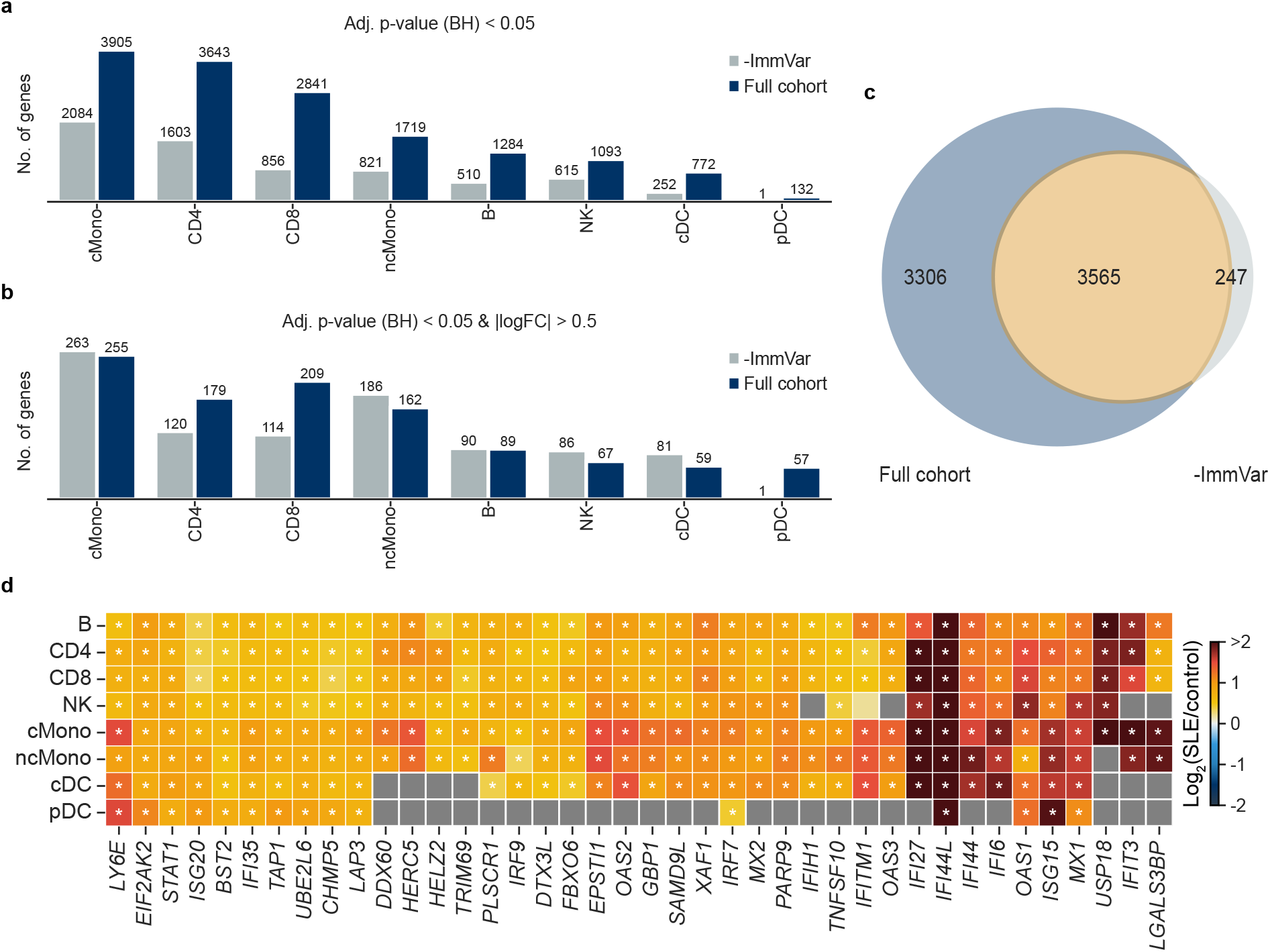
Reanalysis of the Perez et al. (2022) lupus cohort: ImmVar sensitivity and interferon signature. (a) Number of differentially expressed genes (FDR < 0.05) per cell type in the full cohort (261 donors, dark bars) versus the ImmVar-excluded cohort (211 donors, light bars). (b) Same comparison with an additional effect-size filter (| log_2_ FC| > 0.5). (c) Venn diagram of the union of DE genes (FDR < 0.05) across all cell types, showing 3,565 shared, 3,306 detected only in the full cohort, and 247 detected only in the ImmVar-excluded cohort. (d) Heatmap of log_2_ fold change (SLE versus control) for 40 interferon-stimulated genes significant (FDR < 0.05) in at least five of eight cell types. Rows are ordered by biological lineage (lymphoid: B, CD4, CD8, NK; myeloid: cMono, ncMono, cDC, pDC). Columns are clustered by expression similarity (Ward linkage, Euclidean distance). Asterisks denote FDR < 0.05 for that gene–cell type combination. Grey cells indicate genes filtered by filter_by_expr() due to low expression in that cell type. Color scale is clipped at > 2. Cell type abbreviations: B, B cells; CD4, CD4 T cells; CD8, CD8 T cells; NK, natural killer cells; cMono, classical monocytes; ncMono, non-classical monocytes; cDC, conventional dendritic cells; pDC, plasmacytoid dendritic cells.

At a more stringent threshold (FDR < 0.05 and | log_2_ FC| > 0.5), the picture was more nuanced (Figure 3b). T cell populations and pDCs showed substantial gains using the full cohort (CD4: 179 versus 120; CD8: 209 versus 114; pDC: 57 versus 1), while monocyte and other myeloid populations showed comparable or slightly fewer large-effect genes (cMono: 255 versus 263; ncMono: 162 versus 186). This is expected: adding 50 control donors improves the estimation of donor-level variance, producing more accurate *log*_2_FC fold change estimates. Some genes that appeared to have large effects in the underpowered ImmVar-excluded analysis were likely inflated by noisy variance estimates — a form of winner’s curse — and the better-powered full-cohort analysis corrects these toward their true effect sizes, pulling some below the | log_2_ FC| > 0.5 boundary. The genes that remained above the effect-size threshold in the full cohort were therefore more robust in their estimated effects. Plasmacytoid dendritic cells illustrate the most extreme power case: the ImmVar-excluded cohort detected only 1 DE gene at any threshold, while the full cohort detected 132 (FDR only) and 57 (FDR + effect size), reflecting the loss of over half the pDC samples upon ImmVar exclusion.

Across the union of DE genes (FDR < 0.05) from all cell types, 3,565 were shared between the two analyses, 3,306 were detected only in the full cohort, and 247 were detected only in the ImmVar-excluded cohort (Figure 3c). The 3,306 full-cohort-only genes represented signal detectable only when the excluded controls were recovered — genes that were always differentially expressed in SLE but fell below the power threshold without the additional 50 control donors. The 247 genes exclusive to the ImmVar-excluded analysis likely reflect borderline cases that shift across the significance boundary with different sample compositions, consistent with sampling variability rather than true biological differences.

The recovered signal is biologically coherent. Forty ISGs that were significant in at least five of the eight cell types show uniform upregulation across all lineages, with the strongest effects in myeloid populations and moderate but consistent effects in lymphoid populations (Figure 3d). The canonical IFN signature — *IFI44L, MX1, OAS1, ISG15, IFIT3, USP18, IFI6, IFI44, IFI27, OAS3, IFITM1, IRF7*, and *STAT1* — was significant in nearly every cell type tested, demonstrating pan-celltype robustness. Grey cells in the cDC and pDC rows reflect low baseline expression filtered by filter_by_expr() rather than biological absence of the interferon response. This result is not a novel biological finding — the type I interferon signature in SLE is among the most well-established observations in lupus immunology (Baechler et al., 2003; Bennett et al., 2003). Rather, it serves as a validation that dreampy’s mixed-model framework recovers known biology from a dataset where methodological limitations of the original analysis left signal unrecognized. The ImmVar-excluded analysis established a conservative floor of what was detectable even with halved controls, whereas the full-cohort analysis revealed the ceiling gained by modeling batch structure properly and recovering all available samples. The heatmap demonstrates that the recovered signal is biologically coherent, not noise.

## Discussion

dreampy provides a native Python implementation validated against R dreamlet on two independent datasets. Its contribution is not methodological — the underlying statistics are those of limma, edgeR, lme4, lmerTest, pbkrtest, and variancePartition — but architectural: making an established and well-calibrated analytical framework fully accessible within the Python ecosystem where the majority of single-cell preprocessing and analysis now takes place.

dreampy allows users to move seamlessly from AnnData preprocessing through mixed-model DE testing and downstream interpretation without leaving Python, removing the language-switching overhead that can discourage researchers from adopting more appropriate statistical models. The practical value of this accessibility is illustrated by the Perez et al. reanalysis, where a mixed-effects model recovered 50 healthy controls that the original fixed-effects analysis excluded due to batch aliasing (Results).

dreampy complements rather than competes with edgePython (Pachter, 2026), which ports edgeR and extends it with a negative binomial–gamma mixed model following the NEBULA-LN approach. Both packages address the same ecosystem gap — the absence of mature bulk RNA-seq statistical frameworks in Python — but implement fundamentally different statistical pipelines. edgePython operates within the negative binomial GLM framework with cell-level dis-persion shrinkage, while dreampy implements the voom log-linear pipeline with observation-level precision weights and residual variance shrinkage. These frameworks make different modeling assumptions and have different strengths: the voom approach transforms counts to approximate normality and leverages the well-understood machinery of linear (mixed) models, while the NB approach models counts directly. Having both available in Python gives researchers the same breadth of choice that R users have long had between limma-voom and edgeR.

The decomposition of the pipeline into individually callable functions, rather than the two bundled entry points in R dreamlet, offers practical implications beyond convenience. Researchers can inspect intermediate outputs at any stage: examining TMM factors for outlier libraries, verifying that voom weights show the expected mean–variance relationship, checking that mixed-model variance components converge, or comparing eBayes shrinkage before and after moderation. This transparency is especially valuable when adapting the pipeline to non-standard experimental designs where default behavior may not be appropriate. It also facilitates method development: a researcher developing a new variance-stabilizing transformation, for example, can slot it in at the log2cpm() stage and use the downstream pipeline unchanged.

Several directions for future development are planned. Speed optimization — including warm-starting the optimizer from the weight estimation stage, a lean deviance-only fast path, and scratch buffer preallocation — could reduce per-gene fitting overhead, particularly for the cold-start penalty that currently makes dreampy slower than R dreamlet on some datasets. Support for random slopes (currently limited to random intercepts) would enable a broader range of experimental designs. Integration with pathway enrichment tools such as gseapy, while not demonstrated in this paper, is compatible with dreampy’s output format and would complete the analytical loop from counts to biological interpretation. More broadly, dreampy’s modular architecture provides a foundation for extending the pseudobulk mixed-model framework to richer latent variable models for multi-sample single-cell data.

## Limitations

We report several categories of known discrepancy between dreampy and R dreamlet, all documented transparently. dreampy supports random intercepts only; random slopes are not implemented. This covers the most common pseudobulk designs (donor and batch as grouping variables) but excludes designs where the effect of a covariate is expected to vary across groups. We aim to include random slopes in future versions.

dreampy cold-starts the optimizer independently for each gene, while R dreamlet warm-starts from the previous gene’s converged parameters within each parallel worker. On flat or multimodal likelihood surfaces, these different starting points can find different optima, both statistically valid. The cold-start approach is gene-order-independent and fully parallelizable, but it may converge to a different local optimum than R’s warm-start for a small fraction of genes.

When random-effect terms are perfectly collinear, dreampy drops the redundant term before fitting, producing a reduced model. R dreamlet retains the full parameterization. Both approaches yield identical differential expression calls on shared genes, but intermediate quantities (variance components, degrees of freedom) can disagree substantially for the affected assay.

The dreamlet paper describes a two-step voom weighting strategy in which initial Poisson-derived precision weights are used in the first regression pass before fitting the mean–variance trend. However, the R dreamlet code does not implement this two-step procedure — it uses single-pass voom weighting from voomWithDreamWeights(). dreampy matches the R implementation rather than the paper description. Readers comparing our code to the dreamlet’s implementation should note this distinction to avoid mistakenly thinking a step has been omitted.

dreampy defaults to REML for both weight estimation and model fitting, whereas R dreamlet uses ML for weight estimation and REML for model fitting. This is an intentional design choice for consistency and robustness. All cross-language validation in this paper uses R’s ML/REML settings to demonstrate numerical concordance. Users who wish to match R’s behavior exactly can set reml=False in estimate_weights().

Finally, dreampy does not include built-in visualization or pathway enrichment functionality. It is designed as a statistical engine that integrates with the existing Python visualization and enrichment ecosystem rather than reimplementing those tools.

## Methods

### Datasets

**Wells et al. (2025)**. Single-cell RNA-seq of T cells profiling immune aging across donor and tissue site (Wells et al., 2025; CellxGene ID 965008a7-e698-413c-a746-8855a945a7c5). Raw counts were accessed from the.raw slot. Pseudobulk aggregation by donor × tissue site yielded 41 to 153 samples per assay across 13 T cell types (cd3_gd, cd4_cm, cd4_naive, cd4_tem, cd4_temra, cd4_treg, cd4_trm, cd8_cm, cd8_mait, cd8_naive, cd8_tem, cd8_temra, cd8_trm). The DE formula was ∼ *age*_*groups* + (1 | *donor*_*id*) + (1 | *tissue*_*abbrev*), where age_groups is a binary variable (<40 vs >40 years) following the stratification used in the original study.

**Perez et al. (2022)**. Single-cell RNA-seq of peripheral blood mononuclear cells from a lupus cohort (Perez et al., 2022; CellxGene ID 4118e166-34f5-4c1f-9eed-c64b90a3dace). The dataset comprises 1.26 million cells from 261 donors (162 SLE, 99 control) across 4 processing cohorts. Raw counts were accessed from the.raw slot. Pseudobulk aggregation by sample_id (a composite of sample_uuid and Processing_Cohort) × author_cell_type yielded 11 to 336 samples across 10 broad immune cell types (B, T4, T8, NK, cM, ncM, cDC, pDC, PB, Prolif). Figure labels use descriptive names: T4 = CD4 T cells, T8 = CD8 T cells, cM = classical monocytes, ncM = non-classical monocytes, cDC = conventional dendritic cells, pDC = plasmacytoid dendritic cells, PB = plasmablasts, Prolif = proliferating cells. The DE formula was ∼ *sle* + (1 | *donor*_*id*) + (1 | *Processing*_*Cohort*), where sle is a binary indicator derived from the disease column. The donor random effect accounted for 94 technical replicates and 10 longitudinal sample pairs; the processing cohort random effect absorbed batch variation. The biological application (Figure 3) reported results from 8 cell types, excluding plasmablasts (11 samples; collinearity between processing cohort and disease status) and proliferating cells (not a discrete biological subtype).

### Synthetic dataset

Three synthetic designs (fixed effects only, single random effect, crossed random effects) were generated in R with set.seed(11) and exported as CSV reference files. These serve as the CI test suite (109/109 tests passing) and were not used in the preprint figures.

### Gene preprocessing

All real datasets underwent identical gene preprocessing before pseudobulking. Genes from 11 categories of technical artifact were removed by regex-based filtering: canonical histones, immunoglobulin variable/joining segments, ribosomal proteins (cytoplasmic and mitochondrial), mitochondrial-encoded genes, metallothioneins, adaptive TCR variable segments (with exceptions for MAIT, iNKT, and *γδ* markers), hemoglobin and erythroid genes, MTRNR2L pseudogenes, heat shock proteins, and DNAJ co-chaperones. Remaining genes were intersected with a curated protein-coding whitelist derived from GENCODE. The full filter specifications, regex patterns, and exception lists are documented in the repository.

After artifact removal, expression-level filtering (filter_by_expr(), reimplementing the strategy described in Chen et al. (2016) removed genes with insufficient counts for reliable statistical testing, using min_count=5 and min_prop=0.4. Samples with fewer than 20 cells per pseudobulk unit were excluded (min_cells=20), and cell types with fewer than 4 remaining samples were dropped (min_samples=4).

Preprocessing was performed once in Python; both the R and Python validation pipelines consumed the same gene-filtered cell-level input. Pseudobulk aggregation was performed independently within each pipeline.

### Cross-language validation protocol

For each dataset and cell type, R dreamlet and dreampy were run on identical gene-filtered input with matched formulas and REML settings (ML for weight estimation, REML for model fitting — matching R dreamlet’s defaults). A fixed lowess span of 0.5 was used for voom weight estimation in both implementations to isolate validation of the statistical pipeline from smoother-selection variability.

Validation proceeded in three tiers. Tier 1 tests verified exact numerical agreement for quantities computed without optimization: gene filter masks (exact boolean match), TMM normalization factors (rtol=1e-5), and log2 CPM values (rtol=1e-6). Scalar tests verified the eBayes hyperparameters s2_prior and df_prior with rtol=0.005. Tier 2 tests assessed agreement for quantities involving optimization (voom weights, model coefficients, residual standard deviations, unscaled standard errors, variance components, residual degrees of freedom, Satterthwaite degrees of freedom, posterior variances, total degrees of freedom, t-statistics, p-values, and log-odds of differential expression) using a Pearson correlation floor of r ⩾ 0.999, supplemented by a fraction gate requiring that no more than 1–2% of per-gene values exceed a quantity-specific absolute tolerance. For coefficients with per-coefficient output (t-statistics, p-values, degrees of freedom, etc.), each coefficient was tested independently.

This produced 351 total tests across 13 Wells assays (332 passing, 19 failing) and 270 total tests across 10 Perez assays (249 passing, 21 failing). Test counts varied slightly across assays because some quantities were absent when a random-effect term was dropped due to collinearity (e.g., the PB assay’s theta_Processing_Cohort is not tested because dreampy dropped that RE term). For the small number of genes where filter_by_expr() produced different inclusion decisions due to floating-point tie-breaks at the CPM boundary (≈10^−15^ absolute difference), validation was performed on the intersection of retained genes.

### ImmVar sensitivity analysis

The Perez dataset was analyzed in two configurations: the full cohort (261 donors) and an ImmVar-excluded cohort (211 donors) that reproduced the exclusion applied by Perez et al. (2022). Both configurations used the same formula and pipeline settings. Comparison metrics included per-cell-type DE gene counts at FDR < 0.05 and FDR < 0.05 with | log_2_ FC | > 0.5, Venn overlap of DE gene sets across all cell types, and a cross-cell-type heatmap of interferon-stimulated genes significant in at least five of eight cell types.

### Software versions

R analyses used R 4.5.2, dreamlet 1.8.0, variancePartition 1.40.0, edgeR 4.8.0, limma 3.66.0, lme4 1.1.38, lmerTest 3.1.3, pbkrtest 0.5.5, and fANCOVA 0.6.1. Python analyses used Python 3.12.12, dreampy 0.1.0, NumPy 2.3.5, SciPy 1.16.3, pandas 2.3.3, scikit-misc 0.5.2, Py-BOBYQA 1.5.0, and anndata 0.12.6. All packages were installed via conda-forge or pip. The complete Python environment is specified in env.lock.yml in the repository. The complete R sessionInfo() is in validation/R/session_info.txt

## Code Availability

dreampy is available as an open-source Python package at https://github.com/sbwells22/d reampy under the MIT license. The repository includes the full source code, synthetic test suite (109 tests), real-data validation scripts and download instructions for the Wells and Perez datasets, preprocessing scripts, and figure-generating notebooks. Validation metrics and timing benchmarks are included in the repository. Full R dreamlet reference outputs were too large to distribute; a single representative assay (CD4 Treg) is provided to reproduce the figures. Regenerating the complete reference set requires an R installation and the original datasets.

## Acknowledgments

We thank Joe Bell (NVIDIA) for encouraging the use of large language models as a tool for building code bases and for practical guidance in their effective use. We thank Dan Rainbow, and Peter Sims for their encouragement and support throughout this project.

## Reflection

Much of the implementation work in dreampy was carried out with the assistance of Claude Opus 4.5 and 4.6 (Anthropic), which served as a collaborative tool for code generation, debugging, statistical reasoning, and manuscript drafting. The project benefited substantially from this assistance — the translation of a multi-package R statistical pipeline into Python, spanning thousands of lines of numerical code, was completed by a single developer on a timeline that would not have been feasible otherwise.

However, the experience also clarified the current boundaries of what LLMs can reliably contribute to scientific software. This project was, at its core, a translation task: every function had a reference implementation in R, every intermediate value had a ground truth to validate against, and the statistical theory was fully specified in published literature. These reference anchors — technical specifications, original papers, existing codebases, and mathematical derivations — were not optional context but essential inputs without which the LLM’s output would not have been trustworthy. Errors were caught precisely because we could compare against R dreamlet’s output at every pipeline stage; without that scaffold, subtle numerical bugs in variance component estimation or degrees-of-freedom bookkeeping would have been difficult to detect.

We expect that translation projects of this kind will become increasingly tractable as language models improve. De novo projects — work without reference implementations, pre-existing tools, or established theory — present a harder problem. The ease with which an LLM can produce plausible-looking code should not be mistaken for the rigor required to ensure that code is correct. Scientific software development, whether LLM-assisted or not, still demands domain expertise, systematic validation, and healthy skepticism toward any output that has not been independently verified.

## Notes

### Competing Interest Statement

The authors have declared no competing interest.

